# Mapping Functional Protein Neighborhoods in the Mouse Brain

**DOI:** 10.1101/2020.01.26.920447

**Authors:** Benjamin J. Liebeskind, Rebecca L. Young, D. Brent Halling, Richard W. Aldrich, Edward M. Marcotte

**Affiliations:** Department of Molecular Biosciences and Center for Systems and Synthetic Biology, University of Texas, Austin, TX 78712, USA; Department of Integrative Biology and Center for Computational Biology and Bioinformatics, University of Texas, Austin, TX 78712, USA; Department of Neuroscience and Center for Learning and Memory, University of Texas, Austin, TX 78712, USA

## Abstract

New proteomics methods make it possible to determine protein interaction maps at the proteome scale without the need for genetically encoded tags, opening up new organisms and tissue types to investigation. Current molecular and computational methods are oriented towards protein complexes that are soluble, stable, and discrete. However, the mammalian brain, among the most complicated and most heavily studied tissue types, derives many of its unique functions from protein interactions that are neither discrete nor soluble. Proteomics investigations into the global protein interaction landscape of the brain have therefore leveraged non-proteomics datasets to supplement their experiments. Here, we develop a novel, integrative proteomics pipeline and apply it to infer a global map of functional protein neighborhoods in the mouse brain without the aid of external datasets. By leveraging synaptosome enrichment and interactomics methods that target both soluble and insoluble protein fractions, we resolved protein interactions for key neural pathways, including those from refractory subcellular fractions such as the membrane and cytoskeleton. In comparison to external datasets, our observed interactions perform similarly to hand-curated synaptic protein interactions while also suggesting thousands of novel connections. We additionally employed cleavable chemical cross-linkers to detect direct binding partners and provide structural context. Our combined map suggests new protein pathways and novel mechanisms for proteins that underlie neurological diseases, including autism and epilepsy. Our results show that proteomics methods alone are sufficient to determine global interaction maps for proteins that are of broad interest to neuroscience. We anticipate that our map will be used to prioritize new research avenues and will pave the way towards future proteomics techniques that resolve protein interactions at ever greater resolution.

## INTRODUCTION

The complexity of the mammalian brain derives from the connectivity of its constituent cells and the functional modalities of electro-chemical cellular communication. Both aspects of neural complexity, the topology of cellular connections and their functional diversity, are mediated by protein interactions occurring at broad spatial and temporal scales. Due to their common origin, neurons and most other eukaryotic cell types share a large number of discrete macromolecular protein complexes that perform conserved cellular functions. But unlike most cells, neurons also have giant protein interaction webs such as the postsynaptic density, which undergirds the entire postsynaptic membrane and can span up to half a micron (Harris and Weinberg, 2012; Phillips et al., 2001). These webs are not merely structural. Rather, they are dynamic structures whose signaling processes and movements form the basis of complex higher behavioral functions like learning and memory (Boeckers, 2006; Südhof, 2012). Nor are protein interactions the only structural basis for functional groupings in the brain. Lipid rafts cluster ion channels into functional islands in synapses (Dart, 2010; Li et al., 2007), and hyaluronic acid forms the backbone of the neuronal extracellular matrix (ECM), which can regulate neuronal development and activity (Long and Huttner, 2019). These various modes by which proteins cluster into functional neighborhoods in the brain provide unique pathways for behavioral complexity, but also for the emergence of disease. Indeed, many brain diseases result from disruptions in protein interactions (Kuzmanov and Emili, 2013; Li et al., 2015; Malty et al., 2017), though the etiology of complex psychiatric disorders like schizophrenia and autism, and their precise relationship to protein interactions, are only just beginning to be understood (Abraham et al., 2019; Flint and Ideker, 2019; Lyall et al., 2017).

Although novel technologies are allowing neuroscientists to probe the brain’s activity at increasingly acute scales, methods to probe protein interactions and functional neighborhoods on the global scale are lacking. This is due to both inherent and extrinsic difficulties in probing endogenous protein interactions within the brain. First, neural protein complexes are often dynamic, hierarchical, promiscuous, and multi-functional - perhaps more so than in any other tissue type. This creates difficulties for protein interactome determination, which is easier for stable, discrete protein complexes. Second, most of the protein interactions of interest in the brain are of relatively low abundance and associated with membranes, cytoskeletal components, extra-cellular matrix or several of these. Traditional proteomics methods are optimized for abundant, soluble proteins, so these characteristics, especially in combination, make proteomics very challenging.

A variety of new proteomic and computational techniques provide opportunities for inferring global protein interaction networks. Each technique has drawbacks, but by combining them, new insights may be drawn on this very complex problem. One technique involves biochemical separation of native cell extracts followed by shotgun mass-spectrometry to identify the proteins in each fraction. This pipeline allows the determination of protein interactions from co-fractionation profiles of candidate protein pairs, and is therefore called co-fractionation mass-spectrometry (CF-MS). A recent study used CF-MS to measure protein interactions from the mouse brain (Pourhaghighi et al., 2018), resulting in the large-scale first map of soluble protein complexes from the brain. However, one draw-back of standard CF-MS pipelines is their reliance on the solubility of native protein complexes. This is particularly constraining with regards to functional complexes in the brain because many of the protein interactions that are most central to brain function are associated with the membrane, cytoskeleton, or both, and are insoluble in standard native protein preparations, which use gentle, non-ionic detergents. A second barrier to CF-MS of whole brains is the fact that the most important proteins for characteristic brain functions including electrical and chemical signal transmission are not necessarily the most abundant. Synapses, the locus of learning and memory, are a small fraction of the volume of the brain, for instance. This becomes a problem for CF-MS due to the limited sensitivity of even the most cutting-edge mass spectrometers.

To overcome these barriers, we used an integrative proteomics approach to determine a global map of functional protein neighborhoods from the mouse brain (**Figure 1**). Like Pourhaghighi *et al*., we also use CF-MS to create fractionation profiles of soluble complexes from un-enriched brain lysates. However, in order to measure interactions among insoluble proteins, we supplemented conventional CF-MS with two methods employing chemical cross-linking. The first, which we term xCF-MS, involves reversibly cross-linking the samples with formaldehyde and completely solubilizing them with strong denaturants before the CF-MS pipeline (Larance et al., 2016). Formaldehyde is added at a high enough level to stabilize the complexes, but not so high as to completely fix the sample, and the cross-links are denatured by heat before shotgun MS is performed. The second technique is to use MS-cleavable cross-linkers to directly detect protein interactions from the cross-links (XL-MS)(Klykov et al., 2018). In XL-MS, collision-induced dissociation (CID) is used to cleave the linker on the second round of tandem mass-spectrometry (MS^2^), the cleaved pairs are selected based on the known weights of the cross-linker adducts, and a third mass-spectrometry round (MS^3^) is used to fragment and identify each of the two linked peptides, resulting in determination of the two proteins that were linked. Finally, to overcome the limitations imposed by the relatively low-abundance of critical brain proteins, we used cellular fractionation in sucrose gradients to enrich synaptosomes: excised synaptic terminals fused to the postsynaptic membrane of their partner cell.

**Figure 1.**
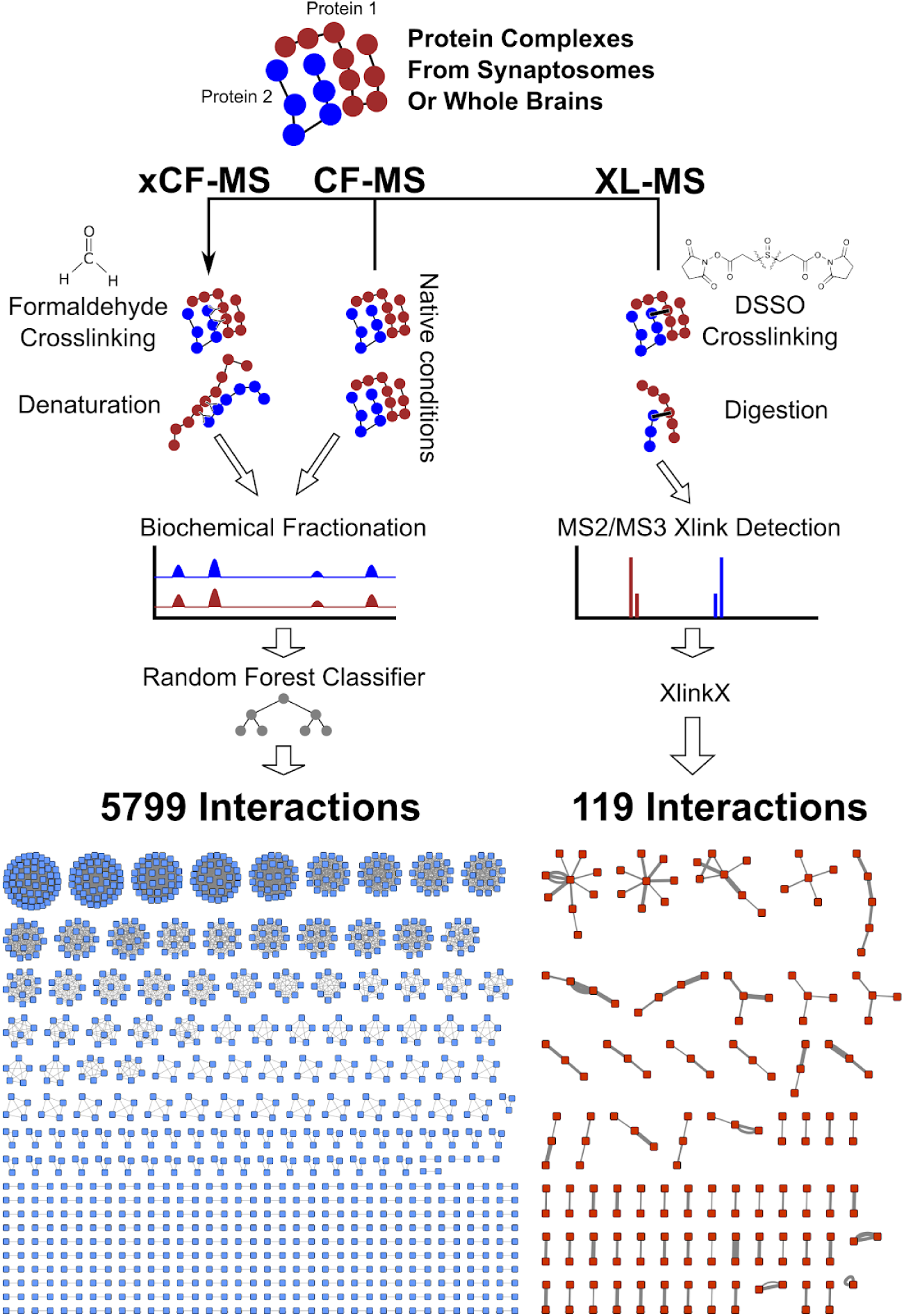
Schematic of integrative proteomics pipeline and resultant interaction maps. Co-fractionation and crosslinking preparations targeting both soluble and insoluble subcellular fractions result in an integrative map of brain protein complexes with attendant structural distance constraints from crosslinking.

This integrative proteomics approach allowed us to improve on current methods and create a map of the functional protein neighborhoods of the mouse brain. Our combined approach allowed us to do so without the addition of external datasets, such as mRNA co-expression, which Pourhaghighi *et al*. used to boost the performance of their method (Pourhaghighi et al., 2018). This map of functional neighborhoods is therefore the first large-scale map of protein complexes in the mammalian brain that relies solely on proteomics of endogenous proteins isolated directly from the mammalian brain.

## METHODS

### Sample preparation

Mouse samples were a gift from the Wallingford and Nishiyama labs at UT Austin. Brains were collected from 3 month old mice, either BL6 or wild-type progeny of WDPCP with a mix of genders, the cerebellums were discarded, and they were immediately frozen in liquid nitrogen before storage at -80°.

For synaptosome preparations, we modified the preparation of Phillips *et al*. (2001). 7 brains were ground in liquid nitrogen with a pre-cooled mortar and pestle to a dry powder. They were then transferred to a 10ml dounce on ice and re-suspended in homogenization buffer (320mM sucrose, 0.1mM CaCl_2_, 1mM MgCl_2_, 0.1mM PMSF). This mixture was homogenized with a Teflon pestle attached to a 4.0 amp Sears Craftsman industrial ⅜ inch drill (model# 315.271490) with 50 down strokes at 2500 rpm. The homogenate was then brought to a final sucrose concentration of 1.25M by adding 2M sucrose in 0.1mM CaCl_2_. After dividing this mixture into ultracentrifuge tubes, we overlaid 1.0M sucrose and equilibrated the tube weights before ultracentrifugation 100,000 x g for 3 hours at 4°. This centrifugation step causes synaptosomes to float to the 1.25/1.0M sucrose boundary, while the bulk of free mitochondria are pelleted together with nuclei on large cell fragments and cytosolic material migrates to the top layer. This synaptosomal band was collected with a 16-gauge needle and either fractionated immediately or stored for later use at -80°.

For xCF-MS samples, we modified the protocol of Larance and colleagues (Larance et al., 2016). They used 6% formaldehyde for 30 minutes to cross-link cell culture samples before denaturation in SDS, but in our hands, these conditions induced aggregation such that samples did not enter diagnostic SDS-PAGE gels. Instead, we cross-linked synaptosome samples in the following way: 1.5ml aliquots of synaptosomes frozen in high sucrose were diluted with 500μl of DPBS and pelleted, and the high sucrose solution removed from the pellet. They were then gently re-suspended with a pipette in 1ml 0.8% formaldehyde DPBS (no more than 1 month old, stored in the dark) and allowed to react for 7 minutes at room temperature before re-pelleting by spinning for 3 minutes at 1,800 x g. The formaldehyde solution was then removed with a pipette and 1ml 100mM Tris added to quench the cross-linking reaction before re-pelleting. For isoelectric focusing, this cross-linked synaptosome pellet was resuspended in 8.4M urea, 2.4M thiourea, 78mM DTT, plus the OFFGel buffer that contains the appropriate ampholytes. This solution was then sonicated on ice with a tip-sonicator at 11% power for 1 minute using 1 second bursts. 1 round of sonication with these conditions was sufficient in this high urea buffer. Cross-linked samples for HPLC were re-suspended in 3M urea, 100mM HFIP and sonicated 3 times with the same conditions, then clarified by centrifugation at 20,000 x g for 10 minutes at 4° and filtered with UltraFree MC 0.45μM spin columns. Urea was kept frozen until use and deionized with mixed bed resin (BioRex RG 501-X8) before making buffers.

All other analyses were performed on single brains. Native samples for CF-MS were ground in liquid nitrogen and homogenized in 150mM NaCl, 0.1mM CaCl_2_, 50mM Tris, 1% n-Dodecyl *β*-D-maltoside (DDM), 0.1mM PMSF, with PhosSTOP tablets, using a glass 2ml dounce, and incubated on ice for 30 minutes. They were then clarified by centrifugation and filtered as above. Brains used for DSSO cross-linking were homogenized similarly, but without DDM.

### Sample fractionation for CF/MS

We performed 9 different fractionation experiments on formaldehyde cross-linked and native samples. For HPLC, we used two kinds of size exclusion columns, a BioSep SEC s4000 and a BioBasic 1000, and two kinds of ion exchange column, a mixed bed PolyLC WAX-WCX, and a concatenation of two anion exchange (PolyWAX) and one cation column (PolyCAT). For isoelectric focusing experiments, we used both pH 3-10 and pH 4-7 strips with 24 wells. Fractionation was performed on either a Thermo Ultimate 3000 HPLC or an Agilent OFFgel system.

After fractionation, formaldehyde cross-linked samples were prepped for mass spectrometry using a protocol modified from Lin and colleagues (Lin et al., 2013) that was optimized for recovery of membrane proteins. To remove urea, IEF fractions were concentrated on 10kD Amicon spin caps, and 10kD Pall plates on a vacuum manifold were used for the same purpose with HPLC fractions. In both cases, the samples were washed once with PBS and then re-suspended in SDC buffer (1% sodium deoxycholate, 50mM NH_4_HCO_3,_ pH 7.5). Samples in native (non-urea) buffers were similarly concentrated but re-suspended in PBS.

All samples were then denatured and reduced in 50% trifluoroethanol and 5mM TCEP by heating in a water-filled heat block at 55° for 45 minutes. They were then cooled to room temperature and alkylated with 15mM iodoacetamide for 30 minutes in the dark before quenching with 7mM DTT. Cross-linked sampled were then diluted to 5% TFE with SDC buffer, and native samples diluted with 50mM Tris, 2mM CaCl_2_, pH 7.5. To these diluted samples we added 1μg of trypsin and digested overnight (16hrs) at 37° after which point the digestion was stopped by acidification to 1% formic acid. Acidification of the cross-linked samples in SDC buffer causes SDC to precipitate, allowing convenient removal of the detergent before mass spectrometry by centrifugation at 2,000 x g. The digestions were then desalted and cleaned by binding to C18 resin, either in spin tips or in plate format, washing with 0.1% formic acid solution, and eluting in 60% acetonitrile, 0.1% formic acid. The elutions were then lyophilized in a vacufuge and re-suspended in 5% acetonitrile, 0.1% formic acid for mass spectrometry.

### Chemical cross-linking

The cross-linking data we report resulted from pooling five different mass spectrometry experiments from two different mouse brain preparations. Both preparations were subjected to subcellular fractionation techniques to simplify the protein sample and to enrich proteins of interest. In particular, we wanted to target proteins that were bound to the cytoskeleton or in lipid rafts that made then insoluble in non-ionic detergents, and hence under-represented in our co-fractionation data. The first preparation targeted proteins from the brain that were not soluble in our 1% DDM preparation of mouse brains used for fractionation (see above). The insoluble pellet was washed twice in DPBS and re-suspended in 300μls of the same before cross-linking. The other preparation targeted synaptosomes and used a subcellular fractionation technique to enrich pre- and postsynaptic complexes (Phillips et al., 2001). Frozen synaptosomes were divided into two aliquots. 2mls of synaptosomes were diluted in 40mls of 1% Triton, 0.1mM CaCl_2_, 20mM MES, pH 6 and incubated on ice for 30mins. This extract was spun down at 10,000 x g for 30 minutes at 4° and the pellet re-suspended 1ml DPBS. The remaining intact synaptosomes (1ml) were diluted in 1ml DPBS, spun down, and re-suspended in 1ml of the same. All the above samples were cross-linked with 1mM DSSO for 1hr with end-to-end rotation at room temperature and quenched with 20mM Tris. After cross-linking, the samples spun down, and both the pellets and the concentrated supernatant were taken through the SDC trypsin digest protocol described above for fractionation.

After cross-linking and tryptic digestion, the first preparation was fractionated on a strong cation exchange (SCX) FPLC column (Thermo HPSP) using a gradient of 5% acetonitrile, 0.1% formic acid pH 2.6 (Buffer C) to Buffer C + 1M KCl. 12 fractions with a high abs 280 signal were kept for analysis. Peptides from the second preparation were fractionated on SCX spin columns (Pierce). First, they were bound to columns and washed with Buffer C, and then sequentially eluted in Buffer C + 100mM KCl, 500mM KCl, and 1M KCl. All SCX fractions were desalted on C18 plates as described above and resuspended in Buffer C for mass spectrometry.

### Mass spectrometry

Mass spectrometry was performed on either a Fusion Lumos Orbitrap or Fusion Orbitrap. Spectra were collected from fractionation experiments using either MS/MS CID or MS/MS HCD with 72 minute gradients.

DSSO cross-links were analyzed using Thermo’s DSSO analysis nodes under either the MS2-MS2 or MS2-MS3 settings. Briefly, MS2-MS3 protocols perform MS/MS-CID, identify the 4 diagnostic peaks of cross-linked peptides, and then submit these peaks to MS3-HCD to be further fragmented. MS2-MS2 performs parallel CID and EThcD MS2 fragmentations, which provide orthogonal spectra for cross-link identification. The first preparation, after SCX fractionation and cleanup, was analyzed under the MS2-MS3 paradigm using a 72-minute gradient. The fractions from the second preparation were analyzed four ways: MS2-MS3 with a 72-minute gradient, MS2-MS3 with a 2-hour gradient (2X), and MS2-MS2 with a 2-hour gradient.

### Proteomic data analysis

Thermo raw spectra files were converted to mzXML format using MSConvert (Adusumilli and Mallick, 2017) and then analyzed using MSBlender (Kwon et al., 2011) integrating analyses from Comet (Eng et al., 2013), X!Tandem (Craig and Beavis, 2004), and MS-GF+ (Kim and Pevzner, 2014). All peptide search engines analyzed the Uniprot mouse reference proteome (UP000000589) with reversed sequences added as decoys and common contaminants. After MSBlender analysis, the number of unique spectral counts for each protein in each fraction was used to create elution profiles for each experiment.

All DSSO cross-linked data was analyzed with XlinkX plugin nodes for Proteome discoverer. We found that the output of this software often found multiple peptides in cross-links that had the same precursor mass and were therefore functionally indistinguishable. We corrected these errors while combining data across multiple samples by first pooling peptide pairs that were found in multiple experiments and taking the highest scoring pair, and then pooling peptide pairs according to calculated precursor mass and leaving only the highest scoring pair. This removed a number of what we deemed to be falsely replicated intermolecular crosslinks.

### Machine learning

From the elution profiles for each experiment, we extracted 5 features for every pair of proteins in the elution: Bray-Curtis distance with column and row max normalization, Pearson’s r and Spearman’s rho with column and row max normalization, and Pearson’s r and Spearman’s rho with column and row sum normalization. Additionally, we concatenated the 9 elution profiles and collected Bray-Curtis, Pearson, and Spearman features with column and row max, and column and row sum normalization. In all these cases, proteins with fewer than 5 total peptide spectral matches in an experiment were removed before normalization. Finally, we added a feature corresponding to the hypergeometric probability of each pair of proteins being found together in a fraction given their background abundances. In total, this yielded 51 features that entered the machine learning pipeline.

For model training and evaluation, we combined three gold-standard datasets: Mouse and rat protein complexes from CORUM (Giurgiu et al., 2019), and mouse complexes from EMBL-EBI’s Complex Portal (Meldal et al., 2015). After combining these datasets, overlapping complexes (with Jaccard coefficient > 0.6) were removed, and the resulting set of 933 complexes was split 60/40% into fully disjoint training and test sets, respectively. These sets of complexes were then resolved into pairs of gene identifiers that were used as positive training and test labels. Negative pairs were generated by drawing random pairs from the input feature set without replacement and keeping pairs that were not in the positive training or test examples. Negative labels were generated in 100-fold excess of the positive training data. After this procedure, the sets used for training consisted of 3383/965192 positive/negative pairs, and the test set contained 2227/426398 positive/negative pairs.

We used TPOT (Olson and Moore, 2016), an automated machine learning algorithm, to guide the machine learning process. TPOT is a genetic algorithm that searches over the space of scikit-learn classifiers, hyperparameters thereof, and pre-processors using cross-validation. Earlier analyses had repeatedly identified the ExtraTrees classifier, a variant of a random forest, as the best classifier in scikit-learn, so we limited TPOT to the ExtraTrees and RadomForest classifiers, while searching over all hyperparameter and pre-processor combinations. TPOT was run on the training data using 50 individuals evolving for 50 generations. After this analysis, TPOT found the best pipeline to be the ExtraTrees classifier, with the following hyperparameters: criterion=“entropy”, max_features=0.1, min_samples_leaf=3, min_samples_split=5, n_estimators=100; and the following preprocessors: ZeroCount() and VarianceThreshold(threshold=0.0005). This pipeline was used to train the model on the entire training set, and this trained model was evaluated on the hold-out test set. We calculated false discovery rate on this hold-out set, found an ExtraTrees score that correlated to 20% FDR, and only considered protein-protein interactions that met this threshold for all downstream analyses.

### Protein and ontology enrichment tests

To assess the contribution of additional co-fractionation mass spectrometry experiments to protein discovery, experiments were subsampled and unique protein ids were counted from each subsampling iteration. For the 14,364 proteins identified, we characterized enrichment patterns including those with high abundance in the synaptosome-enriched samples (756) versus un-enriched samples (11,441) and in the soluble (9,024) versus insoluble (1,138) fractions. Proteins with a one log2 fold difference in abundance are considered enriched. Overlap among Enrichment of Gene Ontology Cellular Component annotations was assessed for both comparisons. Enrichment of GO categories among over- or underrepresented proteins (log2 fold differences in abundance) was calculated using a Mann-Whitney U test (GO_MWU (Wright et al., 2015)). Hierarchical clustering of significant GO categories indicates the intersection gene membership. GO categories connected without branch lengths are subsets of one another. At least 29 genes of a GO category were required to be considered. GO categories containing more than 25% of the gene set were eliminated. GO categories with 90% similarity in gene content were merged to reduce redundancy.

### Network clustering

Protein pairs that met a 20% FDR threshold after the machine learning pipeline were clustered in three ways. First, we used Cluster1 to split the adjacency matrix into discrete clusters to visualize it (**Figure 1**). Second, we used the R package *ggraph* to plot the full FDR-corrected network and compare to external databases. However, we were not satisfied with the interpretability of the Cluster1-defined splits nor with the layout of the entire adjacency matrix. The Cluster1 method may not be best suited to brain PPIs, as many proteins critical for brain function exist not in discrete complexes, but rather in large, macromolecular interconnected structures like the postsynaptic density. Within these structures, proteins exist in smaller organized units. However, plotting the full, highly-connected graph is difficult to interpret. We therefore chose to use a hierarchical clustering approach and to cut the tree at multiple points to examine a protein’s functional neighborhood at varying depths. We used the “cluster_walktrap” method in igraph to create a dendrogram, with 4 steps and a random seed set to 42. We then cut the tree at 9 depths corresponding to 0.9 - 0.1 of the tree’s total height to produce flat clusters.

### Data availability

Our mass spectrometry proteomics data have been deposited to the ProteomeXchange Consortium via the PRIDE partner repository (Perez-Riverol et al., 2019) with the dataset identifiers PXD017110, PXD017111, PXD017136, PXD017137, PXD017138, PXD017175, PXD017176, PXD017178 and PXD017190. On Zenodo (10.5281/zenodo.3565790), we also provide 1.) a spreadsheet of our hierarchically clustered protein pairs with associated annotations and sparklines showing their fractionation profiles 2.) text files of intra- and inter-crosslinks 3.) FDR-corrected PPI pairs inferred by auto-ML 4.) fractionation profiles for each experiment, and 5.) labeled feature matrices for training and test data

## RESULTS

### Native fractionation achieves high coverage of the mouse brain proteome

We sought to directly isolate native multi-protein assemblies directly from mouse brain and semi-purified brain synaptosomes, then to biochemically separate the assemblies sufficiently to identify their components using mass spectrometry. In order to maximally recover brain protein complexes, we employed multiple chromatographic and other biochemical separation methods, in combination with chemical cross-linking, prior to mass spectrometry. In this way, we were able to achieve high coverage of the mouse brain proteome while generally maintaining intact protein assemblies.

Overall, we performed 9 different fractionation experiments on formaldehyde cross-linked and native samples, using 4 distinct fractionation techniques: size-exclusion chromatography (SEC), mixed bed ion-exchange (mbIEX), sequential ion exchange columns (seqIEX), and isoelectric focusing (IEF). Of these 9 fractionation experiments, 4 were enriched synaptosomes analyzed by xCF-MS (IEF and seqIEX), 4 were un-enriched brains analyzed by standard CF-MS (SEC, mbIEX, and seqIEX), and 1 was un-enriched brain analyzed by xCF-MS (mbIEX), leading to the recovery of different subsets of brain proteins (**Figure 2A**). XL-MS studies were performed using the DSSO cross-linker in 5 experiments on un-enriched brain preparations, using both MS2 and MS3-based peptide/protein identification (Liu et al., 2017).

**Figure 2.**
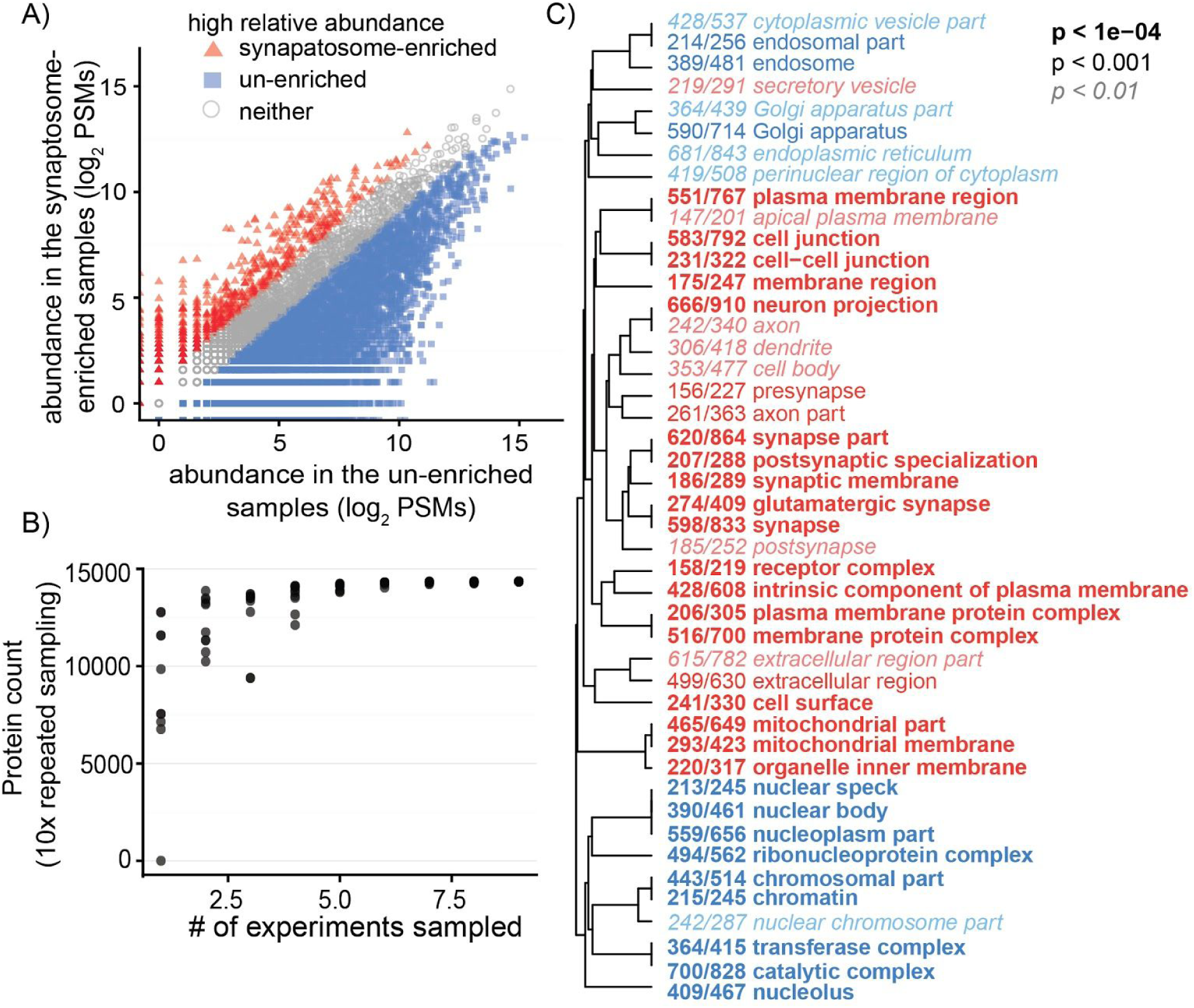
Un-enriched and synaptosome-enriched samples target distinct subsets of the proteome. (A) Enrichment of synaptosomes leads to differential targeting of subsets of the proteome. (B) Saturation analysis suggests that further fractionation experiments with similar parameters would not lead to greater proteome coverage. (C) Gene Ontology (GO) enrichment of proteins abundant in the synaptosome-enriched samples or in the un-enriched samples. The GO categories are clustered by overlapping gene content. The mutual exclusion of synaptosome and brain samples indicates the power of subcellular fractionation to probe less highly expressed proteins.

These combined experiments led to identification of 14,364 proteins that had at least 3 peptide spectral matches (PSMs) in at least one co-fractionation experiment, or a cross-linking score greater than 50. This comprises 91.4% of the mouse brain proteome, as annotated by http://www.mousebrainproteome.com/ (Sharma et al., 2015), and 88.4% of those that are identified as transmembrane proteins by Uniprot (UniProt Consortium, 2019). If we restrict the proteins to those with at least 3 PSMs found in at least 2 experiments, the numbers are 10,973 proteins comprising 83.3% of mouse brain proteins and 76.4% of mouse brain transmembrane proteins. If we apply an even more stringent filter, and require that proteins are found in all soluble fractionations, or all xCF-MS fractionations, or in the cross-linking set, we still recover 8,684 proteins, covering 69.2% of mouse brain proteins and 56.3% of the transmembrane proteins. This represents a substantial increase in coverage over prior brain CF/MS data (Pourhaghighi et al., 2018), and saturation analysis suggests that further experiments using our broad range of conditions are not likely to increase proteome coverage (**Figure 2B**).

We found that proteins in the membrane, cell-junctions, synapses, and mitochondria were highly enriched in the synaptosome samples relative to the un-enriched soluble protein preparations (**Figure 2C**), showing the importance of subcellular fractionation and our xCF-MS pipeline in targeting the characteristic proteome of the brain. We note that the high density of mitochondrial proteins in the synaptosome fractions could be due both to mitochondrial contamination of synaptosome purifications and to synaptosome-associated mitochondria, which are known to interact directly with synaptic machinery (Kang et al., 2008). Soluble fractions were enriched for transcription-translation machinery and other core cellular processes not unique to the brain.

### Machine learning helps determine a high-quality protein interaction map

To determine protein-protein interactions, we considered the co-fractionation and cross-linking experiments separately. XL-MS methods tend to have relatively low coverage due to the fact that only lysine residues within a certain distance of each other will be bound by the cross-linker (Kao et al., 2011), and indeed, the network based only on XL-MS was small (**Figure 1**). However, we note that this method provides structural information whereas co-fractionation does not. We therefore felt that the two methods were best considered as orthogonal information sources.

The network derived from co-fractionation experiments was determined by first extracting standard features (correlation, distance, and hypergeometric co-occurrence features) from each fractionation experiment as well as the merged fractionation profiles, each with two normalization schemes: one with each protein normalized to its PSM sum over fractions, and one with each normalized to its max PSM fraction. This combination of feature extraction methods created 52 starting features for the classifier. We then used automated machine learning methods (auto-ML), as implemented in the Python package TPOT (Olson and Moore, 2016), to calculate protein-protein interaction scores from these features by training it on PPIs from gold-standard databases. Briefly, TPOT uses a genetic algorithm to choose among classifier models and pre-processors by testing each in a cross-validation framework on the training set, and allowing the population of pipelines to “evolve” based on the “fitness” derived from the cross-validation scores. After 50 generations, TPOT chose an ExtraTrees model as the best classifier and two pre-processor steps (see **Methods**). This pipeline was then used to train the model on all of our training data. Performance of the auto-ML model against gold-standard PPIs held out from the training process is shown in **Figure 3B**. We used this recall-precision curve to calculate the false discovery rate (FDR). Proteins with classifier scores corresponding to a 20% FDR or better were used for downstream analyses.

**Figure 3.**
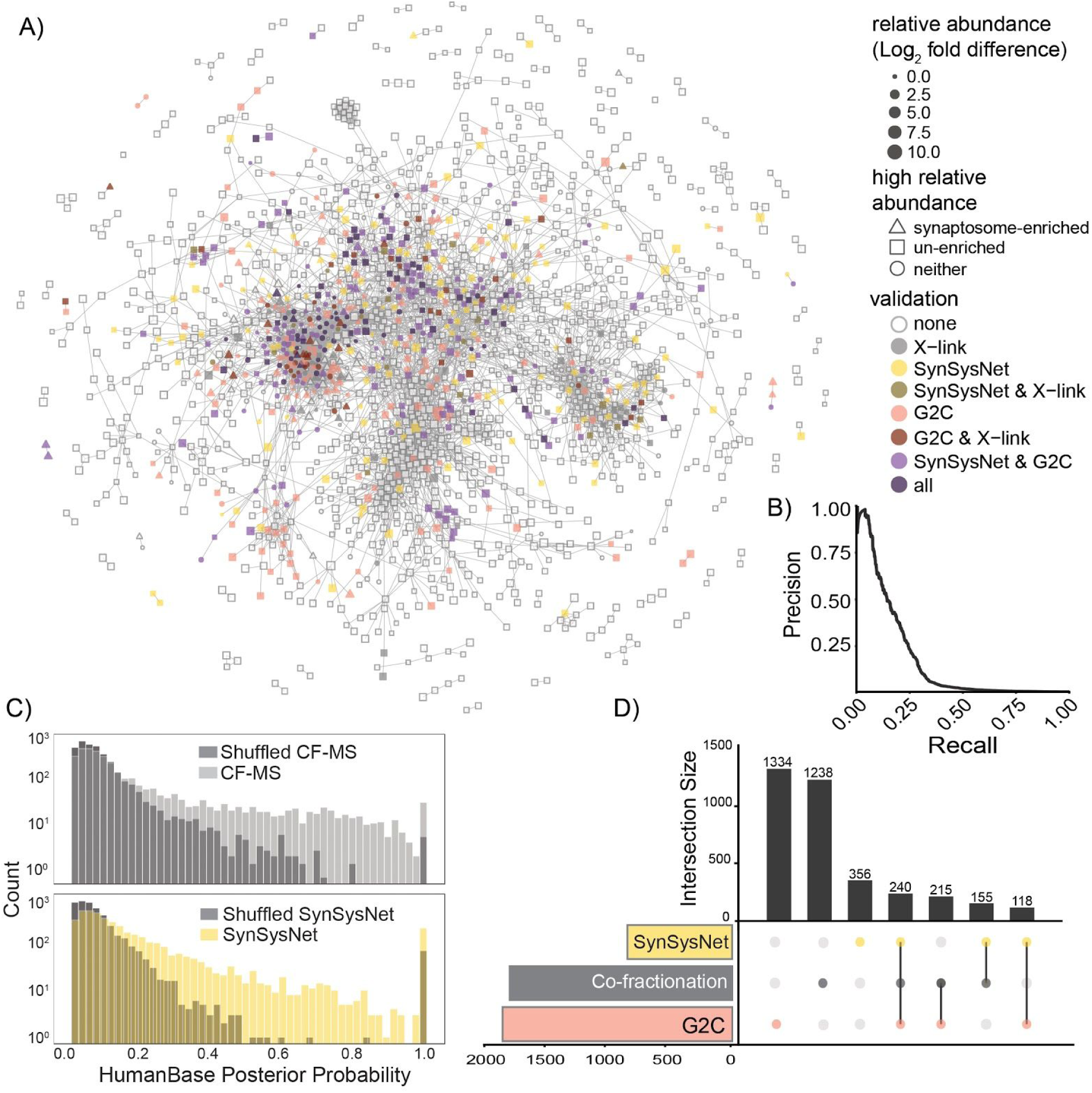
Protein interactions observed by co-fractionation suggest novel protein connections in the synapse. (A) Graph layout of the FDR-corrected adjacency matrix. Parts of the central, highly-connected area of the graph strongly overlap two previous brain protein-interaction databases, SynSysNet and Genes to Cognition (G2C), while other areas are unique. (B) Recall-precision curve on holdout gold-standard protein interaction pairs used to calculate FDR. (C) Measured protein interaction pairs are enriched for higher HumanBase scores over random and are comparable to the hand-curated synapse interaction map SynSysNet. (D) Protein overlap in SynSysNet, G2C, and our interaction set determined from co-fractionation is generally low, but nevertheless highly significant (see **Supplemental Figure 1**). The X-axis to the left shows the number of proteins in each dataset, and the y-axis to the right shows the number that are either unique to each dataset or exist in an intersection of the datasets, denoted by color circles and connecting lines below.

5,799 protein pairs met the 20% FDR threshold and consist of 1,849 unique proteins. We compared the proteins from the well-supported interacting pairs to curated external databases SynSysNet (http://bioinformatics.charite.de/synsys/), containing curated protein pair interactions in the synapse (von Eichborn et al., 2013), and Genes to Cognition (G2C: http://www.g2conline.org/), containing proteins from affinity-purification mass spectrometry experiments (Croning et al., 2009). 455 and 395 of the proteins from the well-supported pairs are contained in G2C and SynSysNet comprising 23% and 45.5% of those databases, respectively. We constructed a protein-protein interaction network with the 1,849 unique proteins as nodes using tidygraph and the ggraph “graphopt” layout (Pedersen, 2018) (**Figure 3A**). Nodes are annotated based on presence in external databases (SynSysNet and G2C), presence in cross-linking mass spectrometry experiments, and their enrichment in the un-enriched versus synaptosome-enriched mass spectrometry experiments. Enrichment is determined as relative abundances in the two regions at +/-1 log2 fold difference. The ExtraTrees classifier score between each protein pairs provide network edge weights. The overlap between our graph and the external datasets G2C and SynSysNet are highly structured. Subsets of the graph correspond strongly to G2C and SynSysNet, while others are unique to our dataset (**Figure 3A**). Overlap in protein sets between the three datasets are generally low (**Figure 3D**).

How well does our large-scale measurement of PPIs correspond to external datasets? We compared our PPIs to the brain-specific interaction scores in HumanBase (Greene et al., 2015), which infers tissue-specific functional interactions by integrating gene-expression and direct interaction data from over 14,000 publications. Our FDR-corrected network was enriched for high HumanBase scores relative to a size-matched random shuffle of the proteins in the FDR-corrected set. The enrichment level was similar to the hand-curated database of synaptic PPIs SynSysNet (von Eichborn et al., 2013)(**Figure 3C**), suggesting that our method produces accuracies that rival the available literature on brain PPIs inferred from standard biochemical methods. Importantly, even though our FDR-corrected PPI set from co-fractionation significantly overlaps SynSysNet (**Supplemental Figure 1**, **Figure 3A**), our set contains many novel proteins (**Figure 3D**) and over 5,500 novel interactions relative to SynSysNet, demonstrating the power of high-throughput proteomics methods for prioritizing new biology for further investigation.

### Co-fractionation broadly recovers neural protein complexes

By combining proteomics methods, we reveal protein interactions that span the range of cellular functions in the brain. Standard CF-MS is well-suited to determining soluble protein complexes (Havugimana et al., 2012; Kirkwood et al., 2013; Pourhaghighi et al., 2018; Wan et al., 2015). Thus, similar to previous studies, we recover conserved, soluble protein complexes such as the proteasome, Cop9 signalosome, and the T-complex/TriC/CCT molecular chaperone (**Supplemental Figure 2**). However, we also recover known complexes from the cytoskeleton, ECM, membrane, pre- and postsynaptic density, and numerous brain-specific pathways. These include known interactions between ion channels, such as BK potassium channels and N-type voltage-gated calcium channels and subunits of ionotropic receptors (Loane et al., 2007).

Our hierarchical clustering method can be used to reveal the nested structures of these interactions. For instance, we find a hierarchy of putative postsynaptic complexes comprising an excitatory module of AMPA receptor subunits, the voltage-gated sodium channel Na_v_1.2, N-cadherin, and the poorly understood protein EFR3b; an inhibitory module comprising the GABA(B) subunits; and a calcium buffering module (**Figure 4**). Though the complex as a whole is undescribed, several members are known to interact, physically or functionally. For instance, N-cadherin and AMPA receptors interact at the cell surface (Nuriya and Huganir, 2006; Saglietti et al., 2007; Silverman et al., 2007), AMPA receptors and Na_v_ channels are both involved in postsynaptic depolarization (Bywalez et al., 2015; Lage-Rupprecht et al., 2019), and EFR3b is a membrane scaffolding protein involved in the synthesis of PIP2, a lipid that regulates AMPA receptor activity (Seebohm et al., 2014) and trafficking (McCartney et al., 2014). In flies, knock-down of the EFR3 homolog “rolling blackout” severely affects neuronal function, but primarily in the pre-synapse (Huang et al., 2004, 2006). In humans, rare variants of EFR3b may be associated with susceptibility to autism, as are variants of Na_v_1.2 (gene name SCN2A), which is also implicated in severe epilepsy and diverse intellectual disabilities (Howell et al., 2015; Shi et al., 2012; Wolff et al., 2017), for example as supported by the FamilieSCN2A Foundation (Scn2a.org). The ATP2 and MCU proteins in the calcium buffering module both act to remove calcium from the postsynaptic intracellular space to the extracellular space and mitochondria, respectively. This serves to maintain synaptic calcium signaling fidelity and prevent toxic calcium buildup (Marland et al., 2016; Pivovarova and Andrews, 2010). Syntaphilin anchors mitochondria, though is thought to do so primarily in axons (Kang et al., 2008). Our hierarchical clustering method therefore provides insight into the functional neighborhoods of proteins involved in synaptic transmission, providing novel insights into the mechanistic basis of disease associated genes like Nav1.2 and EFR3b.

**Figure 4.**
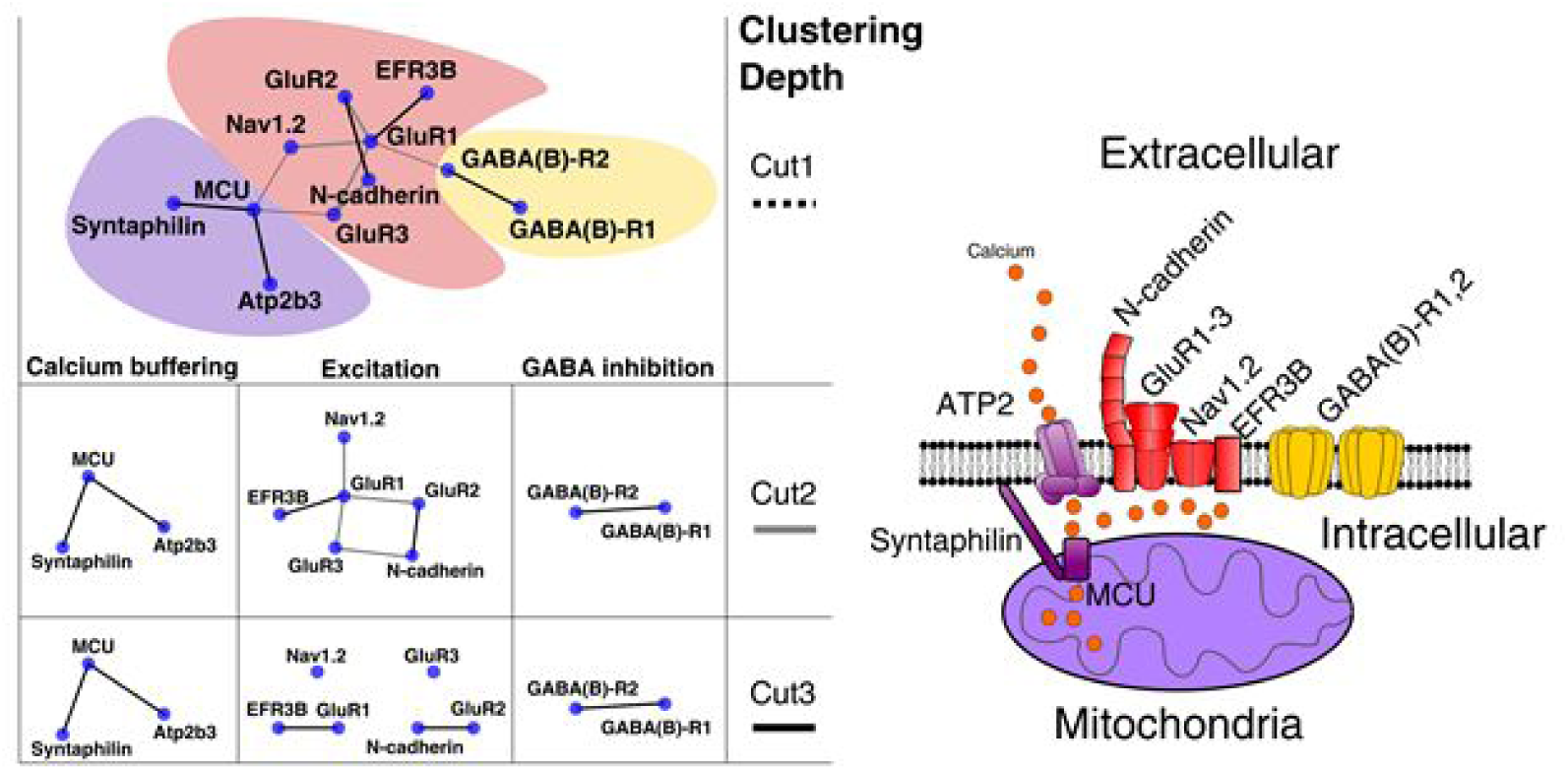
Hierarchical clustering reveals nested functional interactions at the synapse. At three different clustering depths, smaller clusters with higher functional overlap are revealed. The proposed module at the deepest clustering depth is shown at right.

### Cross-linking provides structural support for co-fractionation map

A characteristic of PPIs measured by co-fractionation is that it is not known which interactions are direct and which are transitive (*i.e*., as between subunits of the same complex that do not directly contact one another)(Drew et al., 2017a, 2017b). Though reporting the hierarchical tree of interactions, rather than flat clusters, can help disentangle this issue, XL-MS can provide empirical evidence of direct interactions, and our interaction map determined from XL-MS, though much smaller, still significantly overlaps the co-fractionation map (**Supplemental Figure 1**). For instance, in the pre-synapse, cytoskeletal proteins organize the machinery involved in synaptic exocytosis in a macromolecular structure called the presynaptic particle web (Phillips et al., 2001). Correspondingly, we find a complex composed of cytoskeletal features, mitochondrial proteins, Calcium-calmodulin dependent protein kinase II (CaMKII), and proteins involved directly in synaptic vesicle release. All of these proteins are known to play roles in synaptic transmission, but are they all directly interacting? Most likely not. For instance, the mitochondrial and vesicle fusion elements could be sequentially bound to cytoskeletal elements, and CaMKII, which has both structural and enzymatic functions (Kim et al., 2016), could be bound transiently to each of the other modules. Some of this sub-structure of the complex can be seen in the graph layout (**Figure 5**), and XL-MS supports the direct interactions between CaMKII subunits, ATP synthase subunits, and neurofilament and internexins.

**Figure 5.**
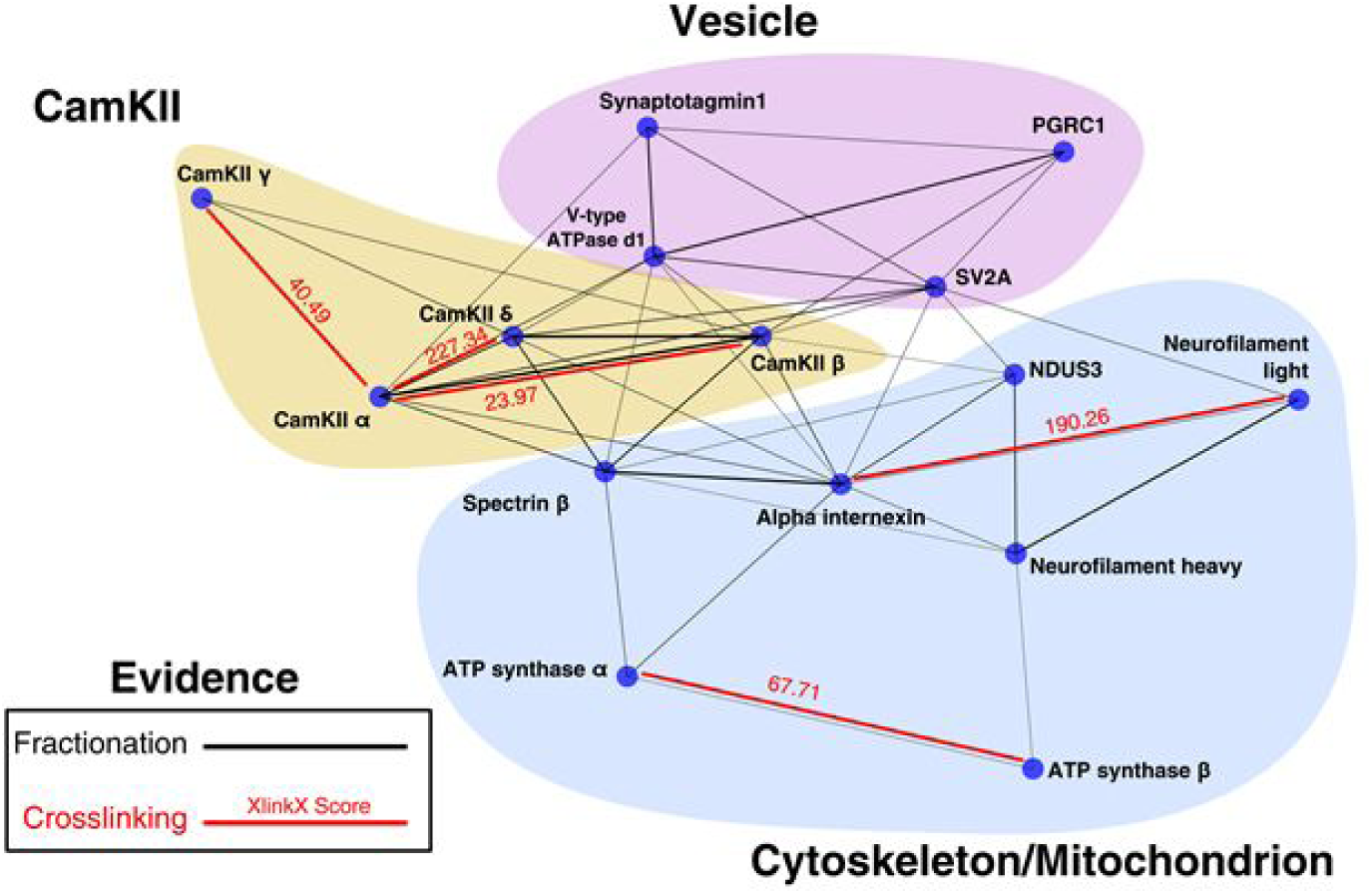
A protein cluster with DSSO cross links annotated. The cross-links recover known interactions between cytoskeletal elements, and the subunits of ATP synthase and CaMKII.

Our XL-MS dataset also provides structural insights. For instance, we recover the known binding relationships of the T-complex subunits *β*, *δ*, *α*, and *γ* (**Figure 6A**). The cross-links also suggest an interaction site between CamKII and mGluRs (**Figure 6B**). In addition to reporting intermolecular cross-links, we found over 400 intramolecular cross-links (chemical cross-links joining two lysines of the same protein) that can inform structural analyses for many proteins critical to brain function, including cell adhesion and cytoskeletal proteins whose structure is not known (**Supplemental Data**).

**Figure 6.**
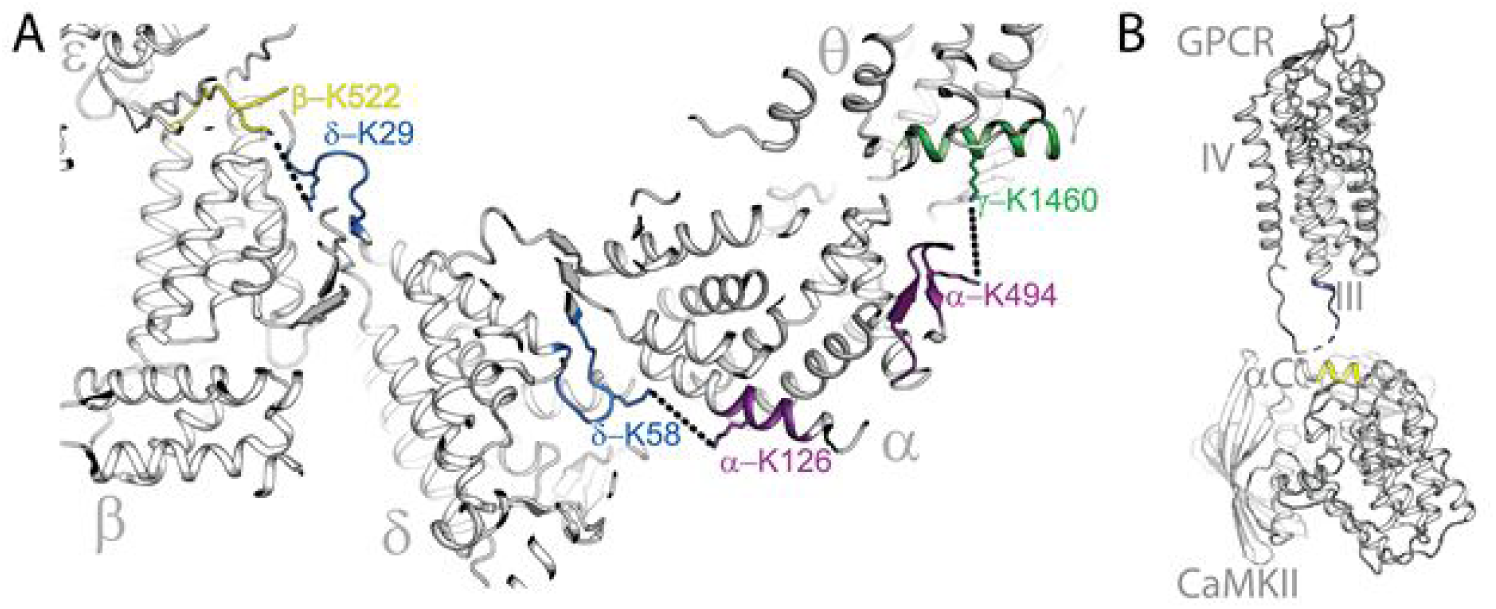
Cross-links reveal structural information about brain protein complexes. Examples include (A) recovering the known subunit structure of the T-complex and (B) suggesting a novel interaction site between CamKII and the metabotropic glutamate receptor (mGluR), a G-protein coupled receptor (GPCR) vital for synaptic transmission and learning and memory.

### Inferred interactions suggest novel mechanisms underlying brain disorders

Protein networks can be used to better understand complex brain disorders. For instance, the molecular mechanisms behind autism spectrum disorders are poorly understood, and many proteins thought to be involved in autism function within large macromolecular structures, making their mechanistic interpretation difficult (Flint and Ideker, 2019). Our study represents one of the few attempts to gain large-scale insights into the connectivity of hard-to-target cellular features like the cytoskeleton, and can therefore shed light on the protein networks that may underlie autism and other brain disorders. We found a number of complexes with subunits implicated in autism (Abrahams et al., 2013)(**Figure 7**). These included groups involved in pathways that are critical for brain function and known to be loci for autism spectrum disorder, like cell adhesion (Stewart, 2015), postsynaptic density (Kaizuka and Takumi, 2018), the cytoskeleton (Lasser et al., 2018). These data provide new mechanistic insights into the protein pathways involved in autism and suggest new candidate proteins for further study.

**Figure 7.**
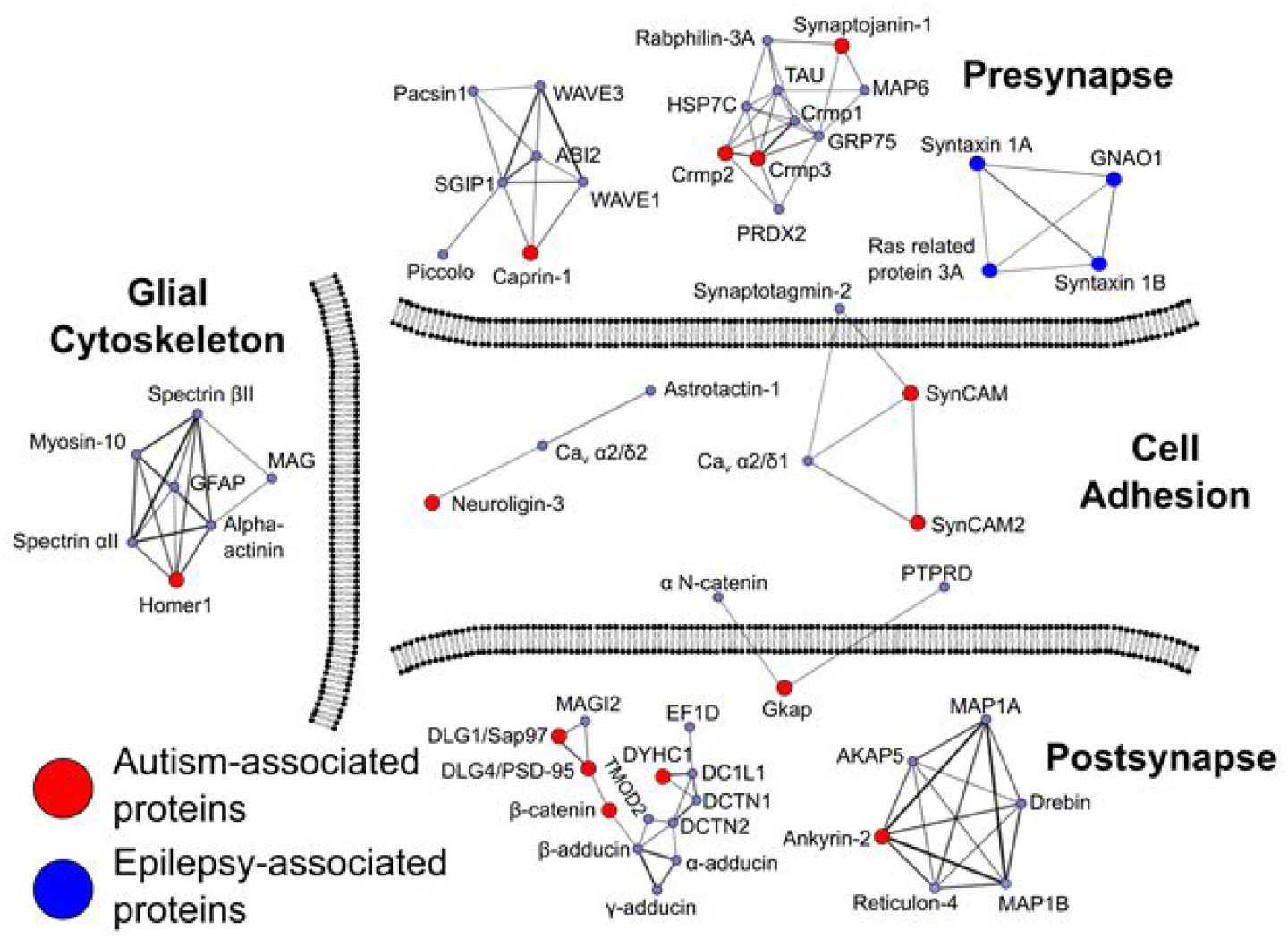
Protein clusters with subunits associated with autism and epilepsy. The clusters contain many proteins known to be involved in synaptic transmission and cell adhesion, and are shown in subcellular locations as annotated in the Uniprot database.

We also found a network comprising proteins involved in synaptic vesicle release and the guanine nucleotide binding protein GNAO1 (**Figure 7**), a highly abundant protein in the brain, mutations in which cause severe epilepsy (Nakamura et al., 2013) or movement disorders (Kulkarni et al., 2016). The mechanistic basis of GNAO1 epilepsy is unknown, but our network suggests that it is caused by disruption of chemical signaling, an inference made stronger by the fact that the other interaction partners, the t-SNAREs syntaxin 1A and B, and Ras-related protein 3A are also implicated in epilepsy (Baghel et al., 2016; Feliciano et al., 2013; Wolking et al., 2019). Pourhaghighi *et al*. 2019 also found an association between GNAO1 and SNAREs (their Figures 2 and 3), though, interestingly, not the same SNARE proteins, and they do not flag GNAO1 as a disease-related protein (their Figure 3). Nonetheless, the general association with SNAREs appears to be independently supported. These findings show the power of global protein interactomics for suggesting novel mechanisms that underlie disease, but call attention to the fact that different proteomics methods and integration of external datasets may alter the specific findings of the study.

## DISCUSSION

Here, we use a novel, integrative proteomics approach to determine the first global protein-interaction map of the brain derived solely from proteomics data. We validated the approach against external databases and found that it performed similarly to hand-curated synaptic protein-interaction databases while also suggesting thousands of novel interactions. We found that our approach, combining soluble and in-soluble fractionation techniques, targets diverse cellular compartments and leads to a network of proteins that span brain functions, including hard-to-target cellular regions like the membrane, cytoskeleton, and ECM. Because many functions unique to the brain occur at the interface of these regions, our approach was able to elucidate functional interactions among proteins of broad interest to neuroscience without the need to incorporate external, non-proteomics datasets. We supplement the co-fractionation interaction network with connections measured by chemical cross-linking, adding angstrom-scale resolution to the protein landscape. Our interaction map suggests novel functional modules and provides mechanistic insights into diverse neuronal diseases.

Global proteomics experiments can help guide and contextualize targeted neuroscience approaches. It is now widely appreciated that the structure and function of neuronal proteins are highly interconnected, often physically. For instance, the original poster-child for a biological transistor, the voltage-gated sodium channel, is now known to reside within large, multi-protein complexes, including structural and signaling elements, as well as other ion channels (Lee et al., 2014; Meadows and Isom, 2005). Indeed, channel isoforms can serve purely as structural elements, rather than channels (Johnson et al., 2018). Furthermore, the evolution of neuronal protein interactions is now known to be more plastic than previously appreciated (Liebeskind et al., 2016), increasing the need for methods such as those we apply here, which are applicable in principle to any organism or cell type. Together, this evidence for an interconnected, pleiotropic, and evolutionarily plastic protein landscape of the brain suggests that increasingly precise, but spatially limited techniques in neuroscience can be usefully complemented by global proteomics methods that can help contextualize more exquisite experimental techniques.

For a novel approach to be maximally useful, it is important to note its caveats as well as its strengths. The most important caveat is that our network is necessarily integrated over the pooled evidence from heterogeneous cell types. Interactions that are more abundant and less transient will be over-represented. For instance, the protein Homer1 is well known as a scaffold in the PSD, but it is actually enriched in glia relative to neurons (Buscemi et al., 2017). Thus our study finds it in a complex with the glia-specific protein GFAP (**Figure 7**). This abundance-weighted global approach should be viewed as an orthogonal line of evidence to traditional, targeted neuroscience approaches that typically start with a protein, cell-type, or brain sub-region of interest.

Our inferred map can also help guide targeted approaches into the mechanistic basis of brain disorders. Our network suggests that the epilepsy-associated protein GNAO1 is a SNARE-interacting protein (**Figure 7**) and that mutations in GNAO1 may therefore cause epilepsy by disrupting synaptic vesicle release. We also implicate a number of proteins in autism through guilt-by-association with proteins already known to be associated with autism (**Figure 7**). These associations are largely found in cell-adhesion and cytoskeletal proteins—regions that are notoriously difficult for proteomics to target, but which we reach with our novel methods.

## CONCLUSIONS

We have introduced a new proteomics pipeline that will be broadly applicable across disciplines, cell-types, and species, as shown here by large-scale analysis of mouse brain protein-protein interactions. It is anticipated that similar combinations of sub-cellular enrichment, integrative proteomics, and machine learning will contribute to a better understanding not only of the mechanisms underlying critical tissue types like the brain, but, by sampling across species and analyzing in a comparative framework (Liebeskind et al., 2019), will also help elucidate the historical process by which these complex molecular machines evolved.

## ACKNOWLEDGEMENTS

The authors gratefully acknowledge Claire McWhite and Kevin Drew for assistance with computational data analysis, John Wallingford, Hiroshi Nishiyama, Janelle Leggere, and Moushumi Dey for providing mouse brains, and Dan Boutz and Ophelia Papoulas for assistance with proteomics methods. We also acknowledge Andrew Horton for identifying and helping correct the XLinkX miscalls described in the Methods. This research was funded by grants from the Welch Foundation (F-1515 to E.M.M.), Army Research Office (W911NF-12-1-0390), and National Institutes of Health (F32 GM112504 to B.J.L. and R35 GM122480 to E.M.M.). R.L.Y. acknowledges funding from the National Science Foundation BEACON Center for the Study of Evolution in Action (DBI-0939454) and a University of Texas CNS Catalyst grant.

---

**Supplemental Figure 1.**
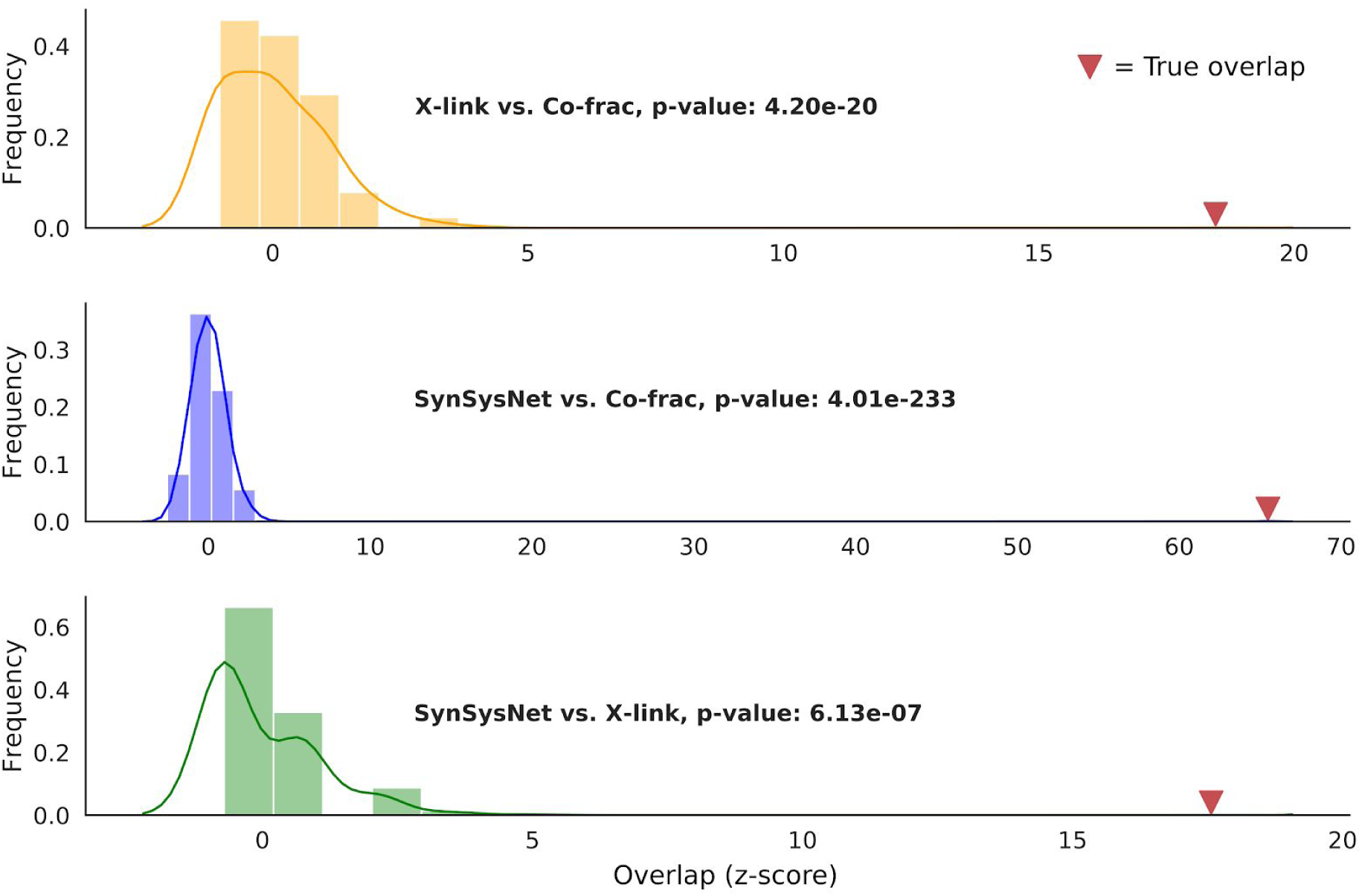
Significance of overlaps between measured and curated literature protein interaction sets. Cross-linking (X-link) and co-fractionation interaction maps significantly overlap each other, and each significantly overlap the hand-curated synaptic protein interaction database SynSysNet. Reported p-values are derived from the hypergeometric test, and distributions show the z-scored overlaps observed from 1,000 random shuffles of proteins within each set, as well as the true overlap.

**Supplemental Figure 2.**
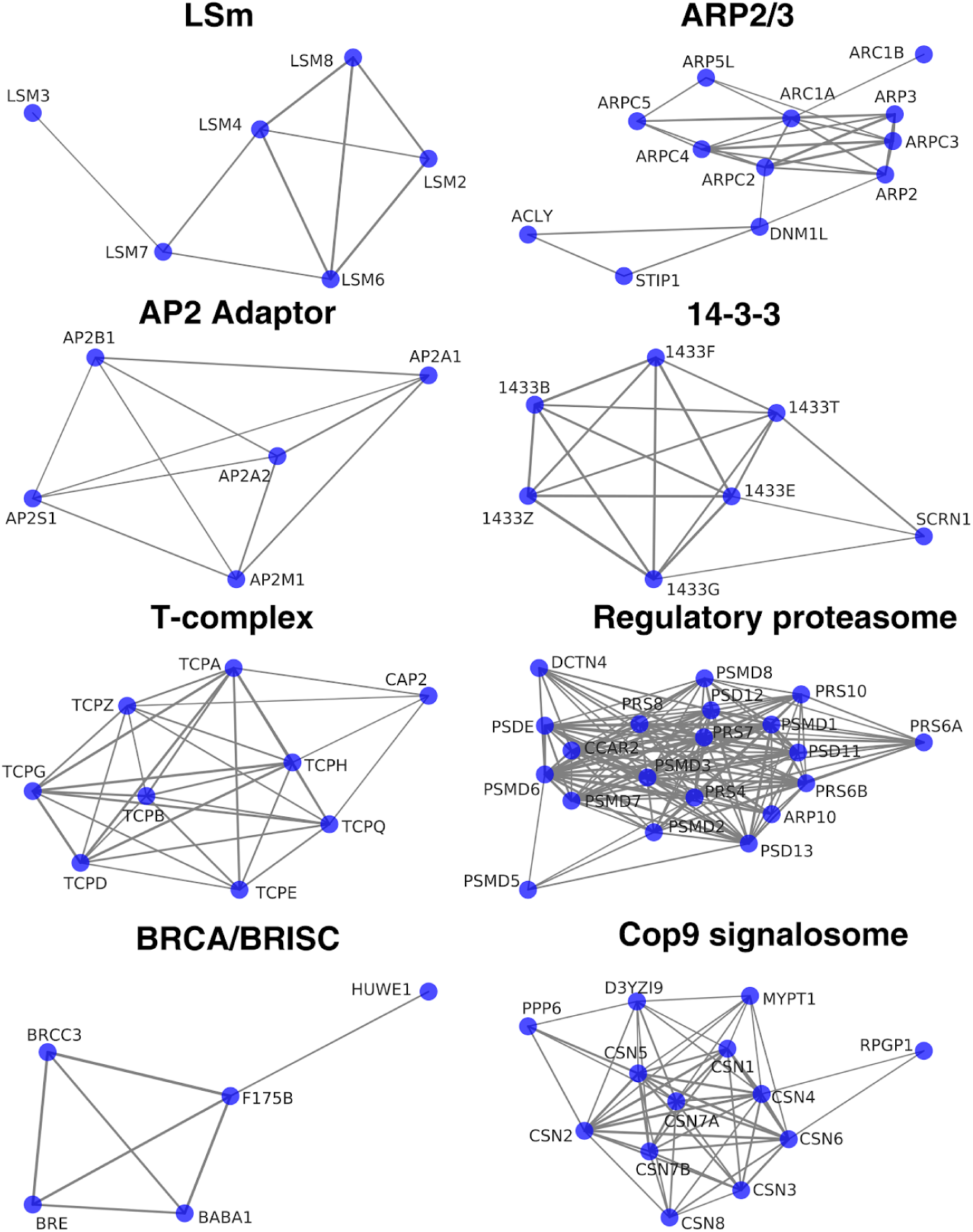
Examples of soluble complexes determined from co-fractionation. The co-fractionation pipeline also recovers known, discrete, soluble complexes that are common to many cell types, as well as neural cells.

## REFERENCES

1. Abraham, J., Szoko, N., and Natowicz, M.R. (2019). Proteomic Investigations of Autism Spectrum Disorder: Past Findings, Current Challenges, and Future Prospects. Adv. Exp. Med. Biol. 1118, 235–252.

2. Abrahams, B.S., Arking, D.E., Campbell, D.B., Mefford, H.C., Morrow, E.M., Weiss, L.A., Menashe, I., Wadkins, T., Banerjee-Basu, S., and Packer, A. (2013). SFARI Gene 2.0: a community-driven knowledgebase for the autism spectrum disorders (ASDs). Mol Autism 4, 36.

3. Adusumilli, R., and Mallick, P. (2017). Data Conversion with ProteoWizard msConvert. Methods Mol. Biol. 1550, 339–368.

4. Baghel, R., Grover, S., Kaur, H., Jajodia, A., Parween, S., Sinha, J., Srivastava, A., Srivastava, A.K., Bala, K., Chandna, P., et al. (2016). Synergistic association of STX1A and VAMP2 with cryptogenic epilepsy in North Indian population. Brain Behav 6, e00490.

5. Boeckers, T.M. (2006). The postsynaptic density. Cell Tissue Res. 326, 409–422.

6. Buscemi, L., Ginet, V., Lopatar, J., Montana, V., Pucci, L., Spagnuolo, P., Zehnder, T., Grubišić, V., Truttman, A., Sala, C., et al. (2017). Homer1 Scaffold Proteins Govern Ca2+ Dynamics in Normal and Reactive Astrocytes. Cereb Cortex 27, 2365–2384.

7. Bywalez, W.G., Patirniche, D., Rupprecht, V., Stemmler, M., Herz, A.V.M., Pálfi, D., Rózsa, B., and Egger, V. (2015). Local postsynaptic voltage-gated sodium channel activation in dendritic spines of olfactory bulb granule cells. Neuron 85, 590–601.

8. Craig, R., and Beavis, R.C. (2004). TANDEM: matching proteins with tandem mass spectra. Bioinformatics 20, 1466–1467.

9. Croning, M.D.R., Marshall, M.C., McLaren, P., Armstrong, J.D., and Grant, S.G.N. (2009). G2Cdb: the Genes to Cognition database. Nucleic Acids Res. 37, D846–851.

10. Dart, C. (2010). Lipid microdomains and the regulation of ion channel function. J. Physiol. (Lond.) 588, 3169–3178.

11. Drew, K., Müller, C.L., Bonneau, R., and Marcotte, E.M. (2017a). Identifying direct contacts between protein complex subunits from their conditional dependence in proteomics datasets. PLoS Comput. Biol. 13, e1005625.

12. Drew, K., Lee, C., Huizar, R.L., Tu, F., Borgeson, B., McWhite, C.D., Ma, Y., Wallingford, J.B., and Marcotte, E.M. (2017b). Integration of over 9,000 mass spectrometry experiments builds a global map of human protein complexes. Mol Syst Biol 13, 932.

13. von Eichborn, J., Dunkel, M., Gohlke, B.O., Preissner, S.C., Hoffmann, M.F., Bauer, J.M.J., Armstrong, J.D., Schaefer, M.H., Andrade-Navarro, M.A., Le Novere, N., et al. (2013). SynSysNet: integration of experimental data on synaptic protein-protein interactions with drug-target relations. Nucleic Acids Res. 41, D834–840.

14. Eng, J.K., Jahan, T.A., and Hoopmann, M.R. (2013). Comet: an open-source MS/MS sequence database search tool. Proteomics 13, 22–24.

15. Feliciano, P., Andrade, R., and Bykhovskaia, M. (2013). Synapsin II and Rab3a cooperate in the regulation of epileptic and synaptic activity in the CA1 region of the hippocampus. J. Neurosci. 33, 18319–18330.

16. Flint, J., and Ideker, T. (2019). The great hairball gambit. PLoS Genet. 15, e1008519.

17. Giurgiu, M., Reinhard, J., Brauner, B., Dunger-Kaltenbach, I., Fobo, G., Frishman, G., Montrone, C., and Ruepp, A. (2019). CORUM: the comprehensive resource of mammalian protein complexes-2019. Nucleic Acids Res. 47, D559–D563.

18. Greene, C.S., Krishnan, A., Wong, A.K., Ricciotti, E., Zelaya, R.A., Himmelstein, D.S., Zhang, R., Hartmann, B.M., Zaslavsky, E., Sealfon, S.C., et al. (2015). Understanding multicellular function and disease with human tissue-specific networks. Nat. Genet. 47, 569–576.

19. Harris, K.M., and Weinberg, R.J. (2012). Ultrastructure of synapses in the mammalian brain. Cold Spring Harb Perspect Biol 4.

20. Havugimana, P.C., Hart, G.T., Nepusz, T., Yang, H., Turinsky, A.L., Li, Z., Wang, P.I., Boutz, D.R., Fong, V., Phanse, S., et al. (2012). A census of human soluble protein complexes. Cell 150, 1068–1081.

21. Howell, K.B., McMahon, J.M., Carvill, G.L., Tambunan, D., Mackay, M.T., Rodriguez-Casero, V., Webster, R., Clark, D., Freeman, J.L., Calvert, S., et al. (2015). SCN2A encephalopathy: A major cause of epilepsy of infancy with migrating focal seizures. Neurology 85, 958–966.

22. Huang, F.-D., Matthies, H.J.G., Speese, S.D., Smith, M.A., and Broadie, K. (2004). Rolling blackout, a newly identified PIP2-DAG pathway lipase required for Drosophila phototransduction. Nat. Neurosci. 7, 1070–1078.

23. Huang, F.-D., Woodruff, E., Mohrmann, R., and Broadie, K. (2006). Rolling blackout is required for synaptic vesicle exocytosis. J. Neurosci. 26, 2369–2379.

24. Johnson, B., Leek, A.N., Solé, L., Maverick, E.E., Levine, T.P., and Tamkun, M.M. (2018). Kv2 potassium channels form endoplasmic reticulum/plasma membrane junctions via interaction with VAPA and VAPB. Proc. Natl. Acad. Sci. U.S.A. 115, E7331–E7340.

25. Kaizuka, T., and Takumi, T. (2018). Postsynaptic density proteins and their involvement in neurodevelopmental disorders. J. Biochem. 163, 447–455.

26. Kang, J.-S., Tian, J.-H., Pan, P.-Y., Zald, P., Li, C., Deng, C., and Sheng, Z.-H. (2008). Docking of axonal mitochondria by syntaphilin controls their mobility and affects short-term facilitation. Cell 132, 137–148.

27. Kao, A., Chiu, C., Vellucci, D., Yang, Y., Patel, V.R., Guan, S., Randall, A., Baldi, P., Rychnovsky, S.D., and Huang, L. (2011). Development of a novel cross-linking strategy for fast and accurate identification of cross-linked peptides of protein complexes. Mol. Cell Proteomics 10, M110.002212.

28. Kim, S., and Pevzner, P.A. (2014). MS-GF+ makes progress towards a universal database search tool for proteomics. Nat Commun 5, 5277.

29. Kim, K., Saneyoshi, T., Hosokawa, T., Okamoto, K., and Hayashi, Y. (2016). Interplay of enzymatic and structural functions of CaMKII in long-term potentiation. J. Neurochem. 139, 959–972.

30. Kirkwood, K.J., Ahmad, Y., Larance, M., and Lamond, A.I. (2013). Characterization of native protein complexes and protein isoform variation using size-fractionation-based quantitative proteomics. Mol Cell Proteomics 12, 3851–3873.

31. Klykov, O., Steigenberger, B., Pektas, S., Fasci, D., Heck, A.J.R., and Scheltema, R.A. (2018). Efficient and robust proteome-wide approaches for cross-linking mass spectrometry. Nat Protoc 13, 2964–2990.

32. Kulkarni, N., Tang, S., Bhardwaj, R., Bernes, S., and Grebe, T.A. (2016). Progressive Movement Disorder in Brothers Carrying a GNAO1 Mutation Responsive to Deep Brain Stimulation. J. Child Neurol. 31, 211–214.

33. Kuzmanov, U., and Emili, A. (2013). Protein-protein interaction networks: probing disease mechanisms using model systems. Genome Med 5, 37.

34. Kwon, T., Choi, H., Vogel, C., Nesvizhskii, A.I., and Marcotte, E.M. (2011). MSblender: A probabilistic approach for integrating peptide identifications from multiple database search engines. J Proteome Res 10, 2949–2958.

35. Lage-Rupprecht, V., Zhou, L., Bianchini, G., Aghvami, S.S., Rózsa, B., Sassoé-Pognetto, M., and Egger, V. (2019). Local reciprocal release of GABA from olfactory bulb granule cell spines: Cooperation of conventional release mechanisms and NMDA receptors. BioRxiv 440198.

36. Larance, M., Kirkwood, K.J., Tinti, M., Brenes Murillo, A., Ferguson, M.A.J., and Lamond, A.I. (2016). Global Membrane Protein Interactome Analysis using In vivo Crosslinking and Mass Spectrometry-based Protein Correlation Profiling. Mol. Cell Proteomics 15, 2476–2490.

37. Lasser, M., Tiber, J., and Lowery, L.A. (2018). The Role of the Microtubule Cytoskeleton in Neurodevelopmental Disorders. Front Cell Neurosci 12, 165.

38. Lee, H.J., Kwon, M.H., Lee, S., Hall, R.A., Yun, C.C., and Choi, I. (2014). Systematic family-wide analysis of sodium bicarbonate cotransporter NBCn1/SLC4A7 interactions with PDZ scaffold proteins. Physiol Rep 2.

39. Li, J., Ma, Z., Shi, M., Malty, R.H., Aoki, H., Minic, Z., Phanse, S., Jin, K., Wall, D.P., Zhang, Z., et al. (2015). Identification of Human Neuronal Protein Complexes Reveals Biochemical Activities and Convergent Mechanisms of Action in Autism Spectrum Disorders. Cell Syst 1, 361–374.

40. Li, X., Serwanski, D.R., Miralles, C.P., Bahr, B.A., and De Blas, A.L. (2007). Two pools of Triton X-100-insoluble GABA(A) receptors are present in the brain, one associated to lipid rafts and another one to the post-synaptic GABAergic complex. J. Neurochem. 102, 1329–1345.

41. Liebeskind, B.J., Hillis, D.M., Zakon, H.H., and Hofmann, H.A. (2016). Complex Homology and the Evolution of Nervous Systems. Trends Ecol. Evol. (Amst.) 31, 127–135.

42. Liebeskind, B.J., Aldrich, R.W., and Marcotte, E.M. (2019). Ancestral reconstruction of protein interaction networks. PLoS Comput. Biol. 15, e1007396.

43. Lin, Y., Liu, H., Liu, Z., Liu, Y., He, Q., Chen, P., Wang, X., and Liang, S. (2013). Development and evaluation of an entirely solution-based combinative sample preparation method for membrane proteomics. Anal. Biochem. 432, 41–48.

44. Liu, F., Lössl, P., Scheltema, R., Viner, R., and Heck, A.J.R. (2017). Optimized fragmentation schemes and data analysis strategies for proteome-wide cross-link identification. Nat Commun 8, 15473.

45. Loane, D.J., Lima, P.A., and Marrion, N.V. (2007). Co-assembly of N-type Ca2+ and BK channels underlies functional coupling in rat brain. J. Cell. Sci. 120, 985–995.

46. Long, K.R., and Huttner, W.B. (2019). How the extracellular matrix shapes neural development. Open Biol 9, 180216.

47. Lyall, K., Croen, L., Daniels, J., Fallin, M.D., Ladd-Acosta, C., Lee, B.K., Park, B.Y., Snyder, N.W., Schendel, D., Volk, H., et al. (2017). The Changing Epidemiology of Autism Spectrum Disorders. Annu Rev Public Health 38, 81–102.

48. Malty, R.H., Aoki, H., Kumar, A., Phanse, S., Amin, S., Zhang, Q., Minic, Z., Goebels, F., Musso, G., Wu, Z., et al. (2017). A Map of Human Mitochondrial Protein Interactions Linked to Neurodegeneration Reveals New Mechanisms of Redox Homeostasis and NF-κB Signaling. Cell Syst 5, 564–577.e12.

49. Marland, J.R.K., Hasel, P., Bonnycastle, K., and Cousin, M.A. (2016). Mitochondrial Calcium Uptake Modulates Synaptic Vesicle Endocytosis in Central Nerve Terminals. J. Biol. Chem. 291, 2080–2086.

50. McCartney, A.J., Zolov, S.N., Kauffman, E.J., Zhang, Y., Strunk, B.S., Weisman, L.S., and Sutton, M.A. (2014). Activity-dependent PI(3,5)P2 synthesis controls AMPA receptor trafficking during synaptic depression. Proc. Natl. Acad. Sci. U.S.A. 111, E4896–4905.

51. Meadows, L.S., and Isom, L.L. (2005). Sodium channels as macromolecular complexes: implications for inherited arrhythmia syndromes. Cardiovasc. Res. 67, 448–458.

52. Meldal, B.H.M., Forner-Martinez, O., Costanzo, M.C., Dana, J., Demeter, J., Dumousseau, M., Dwight, S.S., Gaulton, A., Licata, L., Melidoni, A.N., et al. (2015). The complex portal--an encyclopaedia of macromolecular complexes. Nucleic Acids Res. 43, D479–484.

53. Nakamura, K., Kodera, H., Akita, T., Shiina, M., Kato, M., Hoshino, H., Terashima, H., Osaka, H., Nakamura, S., Tohyama, J., et al. (2013). De Novo mutations in GNAO1, encoding a Gαo subunit of heterotrimeric G proteins, cause epileptic encephalopathy. Am. J. Hum. Genet. 93, 496–505.

54. Nuriya, M., and Huganir, R.L. (2006). Regulation of AMPA receptor trafficking by N-cadherin. J. Neurochem. 97, 652–661.

55. Olson, R.S., and Moore, J.H. (2016). TPOT: A Tree-based Pipeline Optimization Tool for Automating Machine Learning. In JMLR: Workshop and Conference Proceedings, pp. 66–74.

56. Pedersen, T. (2018). Tidygraph: a Tidy API for Graph Manipulation. R Package Version 1.1. 0.

57. Perez-Riverol, Y., Csordas, A., Bai, J., Bernal-Llinares, M., Hewapathirana, S., Kundu, D.J., Inuganti, A., Griss, J., Mayer, G., Eisenacher, M., et al. (2019). The PRIDE database and related tools and resources in 2019: improving support for quantification data. Nucleic Acids Res. 47, D442–D450.

58. Phillips, G.R., Huang, J.K., Wang, Y., Tanaka, H., Shapiro, L., Zhang, W., Shan, W.S., Arndt, K., Frank, M., Gordon, R.E., et al. (2001). The presynaptic particle web: ultrastructure, composition, dissolution, and reconstitution. Neuron 32, 63–77.

59. Pivovarova, N.B., and Andrews, S.B. (2010). Calcium-dependent mitochondrial function and dysfunction in neurons. FEBS J. 277, 3622–3636.

60. Pourhaghighi, R., Ash, P.E.A., Phanse, S., Goebels, F., Malolepsza, E., Tsafou, K., Nathan, A., Chen, S., Zhang, Y., Wierbowski, S.D., et al. (2018). Macromolecular Connectivity Landscape of Mammalian Brain (Rochester, NY: Social Science Research Network).

61. Saglietti, L., Dequidt, C., Kamieniarz, K., Rousset, M.-C., Valnegri, P., Thoumine, O., Beretta, F., Fagni, L., Choquet, D., Sala, C., et al. (2007). Extracellular interactions between GluR2 and N-cadherin in spine regulation. Neuron 54, 461–477.

62. Seebohm, G., Wrobel, E., Pusch, M., Dicks, M., Terhag, J., Matschke, V., Rothenberg, I., Ursu, O.N., Hertel, F., Pott, L., et al. (2014). Structural basis of PI(4,5)P2-dependent regulation of GluA1 by phosphatidylinositol-5-phosphate 4-kinase, type II, alpha (PIP5K2A). Pflugers Arch. 466, 1885–1897.

63. Sharma, K., Schmitt, S., Bergner, C.G., Tyanova, S., Kannaiyan, N., Manrique-Hoyos, N., Kongi, K., Cantuti, L., Hanisch, U.-K., Philips, M.-A., et al. (2015). Cell type- and brain region-resolved mouse brain proteome. Nat. Neurosci. 18, 1819–1831.

64. Shi, X., Yasumoto, S., Kurahashi, H., Nakagawa, E., Fukasawa, T., Uchiya, S., and Hirose, S. (2012). Clinical spectrum of SCN2A mutations. Brain Dev. 34, 541–545.

65. Silverman, J.B., Restituito, S., Lu, W., Lee-Edwards, L., Khatri, L., and Ziff, E.B. (2007). Synaptic anchorage of AMPA receptors by cadherins through neural plakophilin-related arm protein AMPA receptor-binding protein complexes. J. Neurosci. 27, 8505–8516.

66. Stewart, L.T. (2015). Cell adhesion proteins and the pathogenesis of autism spectrum disorders. J. Neurophysiol. 113, 1283–1286.

67. Südhof, T.C. (2012). The presynaptic active zone. Neuron 75, 11–25.

68. UniProt Consortium (2019). UniProt: a worldwide hub of protein knowledge. Nucleic Acids Res. 47, D506–D515.

69. Wan, C., Borgeson, B., Phanse, S., Tu, F., Drew, K., Clark, G., Xiong, X., Kagan, O., Kwan, J., Bezginov, A., et al. (2015). Panorama of ancient metazoan macromolecular complexes. Nature 525, 339–344.

70. Wolff, M., Johannesen, K.M., Hedrich, U.B.S., Masnada, S., Rubboli, G., Gardella, E., Lesca, G., Ville, D., Milh, M., Villard, L., et al. (2017). Genetic and phenotypic heterogeneity suggest therapeutic implications in SCN2A-related disorders. Brain 140, 1316–1336.

71. Wolking, S., May, P., Mei, D., Møller, R.S., Balestrini, S., Helbig, K.L., Altuzarra, C.D., Chatron, N., Kaiwar, C., Stöhr, K., et al. (2019). Clinical spectrum of STX1B-related epileptic disorders. Neurology 92, e1238–e1249.

72. Wright, R.M., Aglyamova, G.V., Meyer, E., and Matz, M.V. (2015). Gene expression associated with white syndromes in a reef building coral, Acropora hyacinthus. BMC Genomics 16, 371.

